# How the Brain Processes Internal and External Sounds: An fMRI Investigation of Auditory Sound Externalization

**DOI:** 10.1101/2025.10.14.679174

**Authors:** Laure Fivel, Jérôme Brunelin, Gaëlle Leroux, Frédéric Haesebaert, Marine Mondino

## Abstract

Auditory externalization refers to the perception of a sound as originating from a source outside the head, as typically occurs in natural listening environments. Conversely, sounds may be perceived inside the head (internalization) when spatial cues are impoverished, as in some headphone-based listening conditions. However, the neural correlates of auditory externalization remain poorly understood. This study investigated differences in brain activation elicited by externalized versus internalized sound sources.

Twenty-nine healthy participants underwent a 3T functional magnetic resonance imaging (fMRI) scan while listening to auditory stimuli presented in three spatialization conditions: two types of externalized sounds (reverberant and anechoic sounds) and one internalized sounds (diotic anechoic sounds).

Whole-brain analyses revealed greater activation for externalized compared to internalized sound sources in the left superior temporal gyrus, including the planum temporale, the cerebellum and the left posterior cingulate gyrus. Internalized sounds elicited greater relative activity in the left inferior temporal gyrus. Direct comparison between the two externalized conditions revealed stronger left superior temporal gyrus activation for reverberant sounds, while anechoic sounds preferentially activated the right middle temporal gyrus.

These findings supported the key role of the planum temporale in auditory externalization and the involvement of higher-order brain regions, suggesting broader networks underpinning the perception of sound location.

## 1. Introduction

The ability to perceive and interpret sounds accurately is a fundamental aspect of human interaction with the world. Auditory perception shapes our daily experiences by enabling us to navigate in our environment, detect and respond to potential threats and communicate effectively. One of the key elements of auditory perception is sound source localization, the process by which we determine the spatial origin of sounds in the environment. In our daily livings, we have to process sounds originating from external objects in the environment. These sounds are typically perceived as originating from a source outside the head, i.e., externalized. This critical aspect of auditory perception is referred to as *externalization* (Best et al. 2020).

Auditory externalization relies on the integration of multiple spatial cues. These include binaural differences in timing (interaural time differences) and intensity (interaural level differences), spectral cues resulting from the filtering effects of the listener’s body (e.g., torso, ears and pinnae), and reverberation cues resulting from the interaction of sound waves with surrounding surfaces (Leclère et al. 2019). When one or more of these cues are absent or degraded, the perception of externalization is weakened. This typically occurs in headphone listening, where sounds presented diotically, i.e., identically to both ears, lack interaural and reverberant information, leading to the perception that sounds originates within the head, i.e., the sound source is internalized. This internalization of sound, although acoustically artificial, provides a unique experimental handle to study the mechanisms that support the construction of spatial auditory representations. Indeed, the perceptual switch between internalized and externalized sounds offers a powerful model for investigating how the brain integrates sensory and contextual cues to attribute spatial origin to auditory events.

Although this perceptual attribution of the sound source as externalized is a fundamental component of auditory scene analysis, its neural basis remains largely underexplored. Most imaging studies have focused on spatial hearing more generally or examined internalized and externalized sounds separately (Krumbholz et al. 2005a; Deouell et al. 2007). To our knowledge, only two functional magnetic resonance imaging (fMRI) studies, albeit with small samples, have directly contrasted externalized versus internalized sound sources (Hunter et al. 2003; Callan et al. 2013). Both reported greater activation in the left planum temporale (PT), a region within the posterior-superior temporal gyrus (STG) for externalized compared to internalized sound sources. Interestingly, Krumbholz et al., (2005a) observed reduced activity of the STG for internalized sound sources compared to monaural spatial sounds, suggesting that internalized sources did not engage the location processing networks in the brain. However, findings regarding other brain regions have been inconsistent. While the STG is central to spatial cue processing, externalized sounds may additionally engage other higher-order regions involved in spatial hearing. Indeed, spatial hearing is thought to rely on a dorsal “where” pathway that projects to regions such as the inferior parietal lobule and the dorsolateral prefrontal cortex (Maeder et al. 2001; Alain et al. 2001; Rauschecker 2018; van der Heijden et al. 2019).

These findings highlight the complexity of externalization process, but the underlying brain networks remain only partially characterized. Importantly, previous neuroimaging studies used externalized stimuli such as sounds convolved with anechoic head-related transfer functions, (HRTF) that lacked reverberation cues, which may have resulted in a weaker sense of externalization and, consequently, reduced neural activation (Leclère et al. 2019). In this context, the present study aimed to further elucidate the neural correlates of auditory externalization by using fMRI in a larger sample than previous studies. For this purpose, we processed human vocal sounds to achieve three levels of externalization when presented through headphones: diotic anechoic sounds, perceived as originating inside the head (internalized); sounds convolved with anechoic HRTFs, perceived as externalized; and sounds convolved with binaural room impulse responses (BRIR), which is supposed to provide stronger externalization due to the addition of simulated reverberation (Leclère et al. 2019).

We hypothesized that activation in the left PT will vary depending on the degree of externalization, being strongest for externalized compared to internalized sound sources. Moreover, as reverberant sounds are more likely than anechoic sounds to be perceived as externalized (Leclère et al. 2019), we expected stronger activation of the left PT for BRIR-convolved compared to HRTF-convolved sounds. In addition, we predicted that more externalized sounds would elicit stronger activation in higher-order brain regions of the dorsal auditory pathway.

## 2. Methods

### 2.1. Participants

Thirty healthy adults were recruited in the study via social networks and university mailing list between 16 January 2024 and 1 July 2024. All participants provided written informed consent. They were right-handed, had no medical treatment (with the exception of oral contraception for women), no current or past personal neurological or psychiatric disorders or in their first-degree relatives as assessed using a structured clinical interview, and no contraindications for MRI, including pregnancy, surgery and metallic or electronic implants. None of participants reported any hearing impairment, and regular practice of a musical instrument avoided the potential confounding effects of musical training on auditory perception (Micheyl et al. 2006). None had consumed alcohol in the 24 hours or coffee in the 12 hours before the MRI session. One participant did not complete the MRI session due to claustrophobia leading to perform the analyses on twenty-nine participants (14 women / 15 men, mean age ± standard deviation = 22.48 ± 2.34 years, mean years of education = 15.17 ± 2.24).

This study was performed in line with the principles of the Declaration of Helsinki. Approval was granted by the Ethics Committee *Comité de Protection des Personnes Est II, France* on May 30, 2022. The study is the neuroimaging part of a larger study preregistered on June 16, 2023 (NCT05936307;https://clinicaltrials.gov/study/NCT05936307).

### 2.2. Stimuli and procedure

Eight neutral vocalizations of the vowel /a/ pronounced by four males and four females’ actors were selected from the universal Montreal Affective Voices battery (Belin et al. 2008). The binaural stimuli were processed to generate three different levels of externalization (Leclère et al. 2019): 1-diotic stimuli (i.e., binaural stimuli presented simultaneously to both ears), assumed to be internalized and perceived inside the head; 2-stimuli filtered by a binaural room impulse response (BRIR) measured by Hummersone et al. 2010 in a theater (room C), which simulates a reverberant environment such as a room and assumed to be externalized and perceived outside the head, even with headphones. 3-stimuli filtered by an anechoic head-related transfer function (HRTF) obtained from anechoic measurements, which simulates the body characteristic effect on sound waves (i.e., body, head, pinnae) and creates an intermediate perception of perceived externalization (i.e., outside the head). The HRTF- and BRIR-filtered voices were delivered from one of four azimuths (-90°, -60°, 60°, 90°; angles <0° were in the left hemifield) to induce variability in source localization on the horizontal plane compared to the frontal position of diotic stimuli. Stimuli were standardized to be delivered with the same duration (1 s) and equalized in volume level.

Participants completed 6 runs of 60 trials (20 diotic, 20 HRTF, 20 BRIR) generating a total of 360 trials. Each run lasted approximately 6 min 30 s. The order of trials was randomized within each run using optimizeX (http://www.bobspunt.com/easy-optimize-x/; further details are available in Supplementary Material 1). The stimuli were delivered through MR-compatible headphones of the OptoAcoustic system of the fMRI at a comfortable listening level. An adjustment of the sound volume was made before the first run to ensure participants could hear the stimuli clearly above the MRI noise. During each trial, an active noise cancelling was performed with OptoAcoustic system and participants listened to the stimulus with their eyes closed. Prior to the fMRI session, participants completed a practice trial to ensure they could reliably perceive the sound sources. The runs were programmed using PsychoPy3 (v2020.2.10).

### 2.3. MRI data acquisition and preprocessing

Imaging data were acquired on a 3T Siemens Magnetom Prisma scanner using a 64-channel head coil at the CERMEP imaging center (Bron, France). A 3D T1-weighted anatomical sequence covering the whole brain volume was collected with the following parameters: 240 slices, TR/TE/TI = 2500/2.22/1000 ms, slice thickness = 0.8 mm, flip angle = 8°, field of view = 256 mm, acquisition time = 6 min 54 s. For each run, fMRI data were acquired using multiband echo-planar imaging (EPI) sequences with the following parameters: 63 axial slices, slice thickness = 2.20 mm, multiband factor = 3, TR/TE = 1500/30 ms, flip angle = 80°, field of view = 210 mm, acquisition time = 6 min 40 s. A total of 1567 volumes were obtained for each participant (between 260 and 264 volumes per run).

Functional images were preprocessed using fMRIPrep 23.2.0 (for details, see Supplementary Material 2). The slice timing was corrected. The images were realigned, resampled to 2-mm cubic voxels and spatially normalized into the Montreal Neurological Institute (MNI) stereotaxic space. Quality checks were carried out using fMRIPrep and MRIqc 23.1.0 outputs.Further preprocessing steps were carried out using SPM12.v7771, including spatial smoothing with an 8-mm full width at half maximum isotropic Gaussian kernel.

### 2.4. fMRI data analysis

Preprocessed fMRI data were analyzed using SPM12.v7771. For each individual, a general linear model was estimated with three regressors of interest (BRIR, HRTF, diotic). Fifteen regressors from previous preprocessing steps were added as regressors of no interest to improve the denoising and account for artifacts such as head motion. These included the 6 motion regressors and their derivatives, as well as the global signal, the white matter and cerebrospinal fluid (CSF) signals. A high-pass filter of 128 s was applied to remove low-frequency noise. All covariates were convolved with the canonical hemodynamic response function and within-subject contrasts were generated for the contrasts of interest: [(HRTF + BRIR) > diotic], [diotic > (HRTF + BRIR)], [BRIR > HRTF] and [HRTF > BRIR]. One-sample t-tests were conducted to analyze the contrasts in blood oxygenation level-dependent (BOLD) signal intensity. Results were thresholded at a voxel-level threshold of p < 0.001 (uncorrected), with a subsequent cluster-level family-wise error (FWE) correction for multiple comparisons at p< 0.05. Results surviving voxel-wise FWE correction (*p* < 0.05) are also reported. Functional activations were anatomically labeled using the automated anatomical labeling 3 (AAL3) atlas implemented in SPM (Rolls et al. 2020).

## 3. Results

The results of whole brain analyses are detailed in Table 1.

**Table 1.**
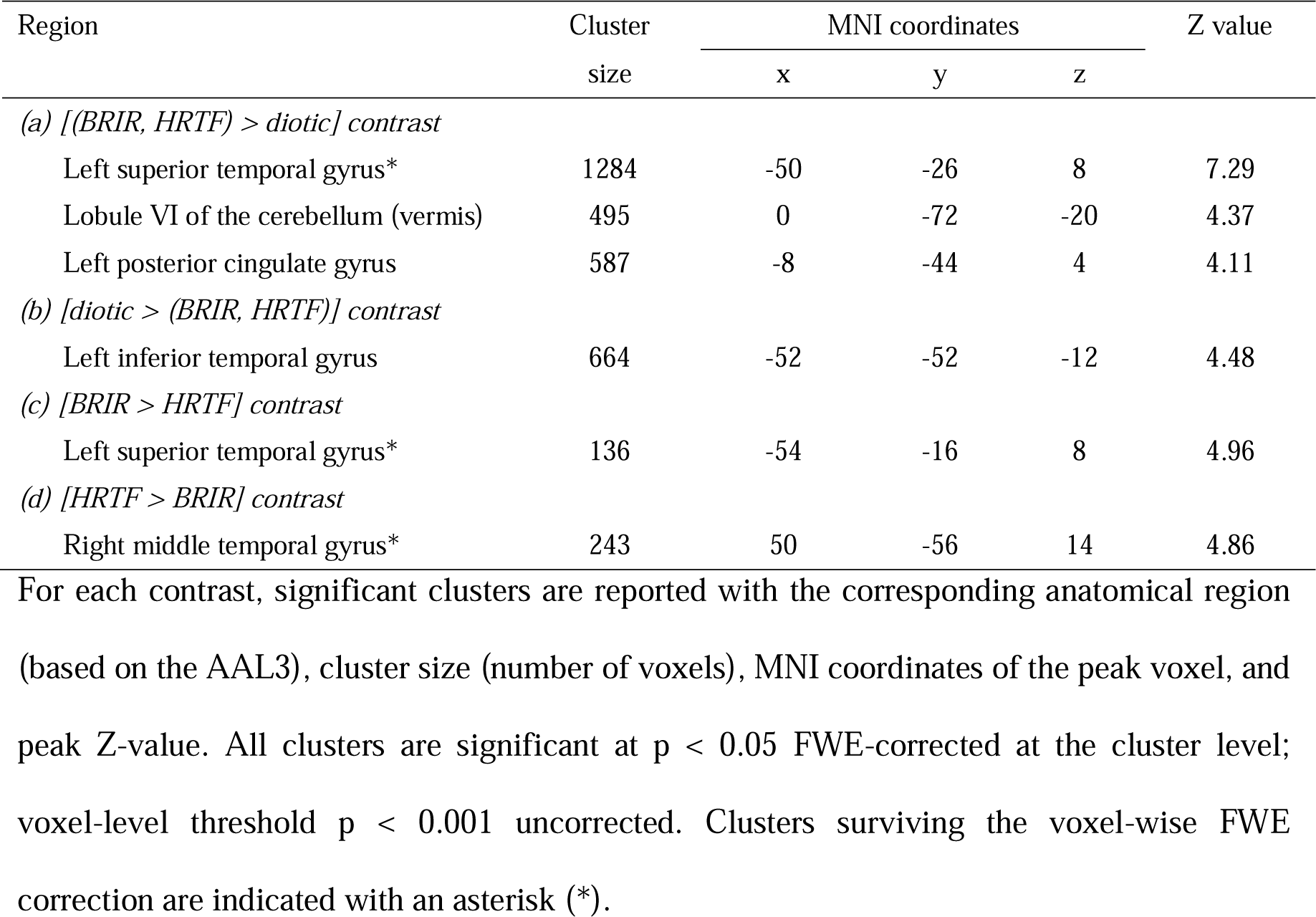
Brain regions with significant differences for each contrast of externalized and internalized sound sources.

| Region | Cluster<br>size | MNI coordinates |  |  | Z value |
| --- | --- | --- | --- | --- | --- |
|  |  | x | y | z |  |
| *(a) [(BRIR, HRTF) > diotic] contrast* |  |  |  |  |  |
| Left superior temporal gyrus* | 1284 | -50 | -26 | 8 | 7.29 |
| Lobule VI of the cerebellum (vermis) | 495 | 0 | -72 | -20 | 4.37 |
| Left posterior cingulate gyrus | 587 | -8 | -44 | 4 | 4.11 |
| *(b) [diotic > (BRIR, HRTF)] contrast* |  |  |  |  |  |
| Left inferior temporal gyrus | 664 | -52 | -52 | -12 | 4.48 |
| *(c) [BRIR > HRTF] contrast* |  |  |  |  |  |
| Left superior temporal gyrus* | 136 | -54 | -16 | 8 | 4.96 |
| *(d) [HRTF > BRIR] contrast* |  |  |  |  |  |
| Right middle temporal gyrus* | 243 | 50 | -56 | 14 | 4.86 |

For each contrast, significant clusters are reported with the corresponding anatomical region (based on the AAL3), cluster size (number of voxels), MNI coordinates of the peak voxel, and peak Z-value. All clusters are significant at $p < 0.05$ FWE-corrected at the cluster level; voxel-level threshold $p < 0.001$ uncorrected. Clusters surviving the voxel-wise FWE correction are indicated with an asterisk (\*).

### 3.1. Outside vs. Inside the head

The contrast [(BRIR + HRTF) > diotic] revealed three significant clusters (Figure 1A): one in the posterior portion of the left superior temporal gyrus corresponding to the planum temporale (MNI [-50, -26, 8], z = 7.29; k = 1284; p_cluster-level FWE_ < 0.001), one in the cerebellum, specifically lobule VI of the vermis ([0, -72, -20]; z = 4.37; k = 495; p_cluster-level FWE_ < 0.001), and one in the left posterior cingulate gyrus ([-8, -44, 4]; z = 4.11; k = 587). Only the left planum temporale cluster survived the voxel-wise FWE correction (p < 0.05).The contrast [diotic > (BRIR + HRTF)] revealed a single significant cluster, in the left inferior temporal gyrus (ITG; [-52, -52, -12]; z = 4.48; k = 664; p_cluster-level FWE_ < 0.001; Figure 1B). This cluster also overlaps with the middle temporal gyrus (peak [-60, -36, -2]).

**Fig. 1.**
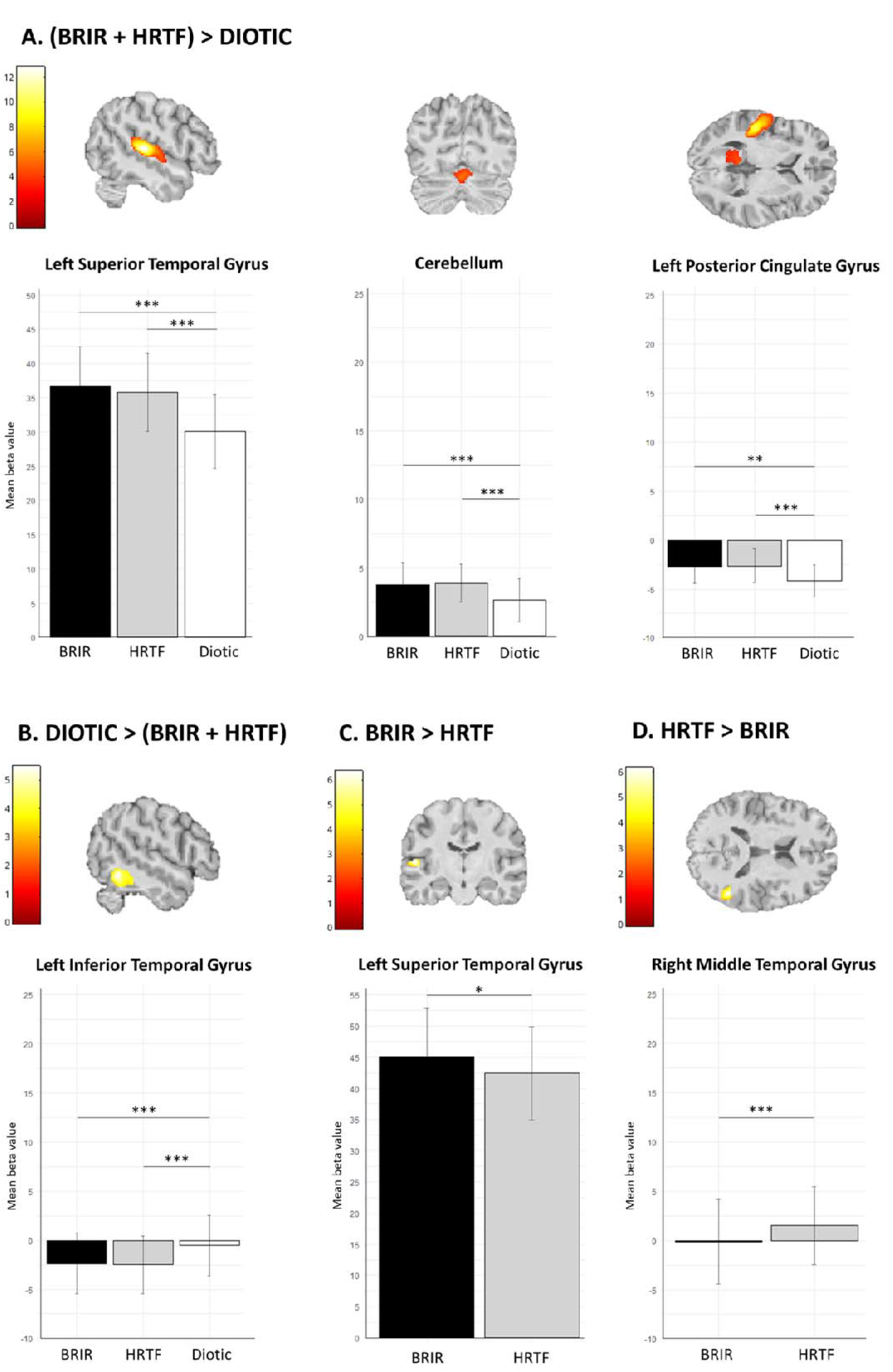
Brain regions showing significant activations for the contrasts. **A)** externalized sounds (sounds filtered by Head Related Transfer Function - HRTF and Binaural Room Impulse Response - BRIR) minus internalized sounds (diotic), **B)** internalized sounds (diotic) minus externalized sounds (HRTF and BRIR), **C)** reverberant sounds (BRIR) minus anechoic sounds (HRTF) and **D)** anechoic sounds (HRTF) minus reverberant sounds (BRIR). *t*-maps are thresholded at *p*_unc_ < 0.001 and cluster-level FWE correction at *p*_FWE_ < 0.05. Images are displayed on the T1 anatomical brain of a single-subject, in axial, coronal and sagittal views. Beta estimates of parametric modulation in the significant clusters are displayed as the mean and standard deviation. \*\**\*p<0.001.* \**\*p<0.01. *p<0.05.*

### 3.2. BRIR vs. HRTF sounds

The contrast [BRIR > HRTF] revealed a significant cluster in the left superior temporal gyrus, with a peak at [-54, -16, 8] (z = 4.96, k = 136; p_cluster-level FWE_ = 0.049; Figure 1C).The reverse contrast, [HRTF > BRIR] revealed a significant cluster in the right middle temporal gyrus, with a peak at MNI coordinates [50, -56, 14] (z = 4.6, k = 243; p_cluster-level FWE_ = 0.005, Figure 1D). Both clusters survived the voxel-wise FWE correction.

A summary of all the contrasts is depicted in Figure 2.

**Fig. 2.**
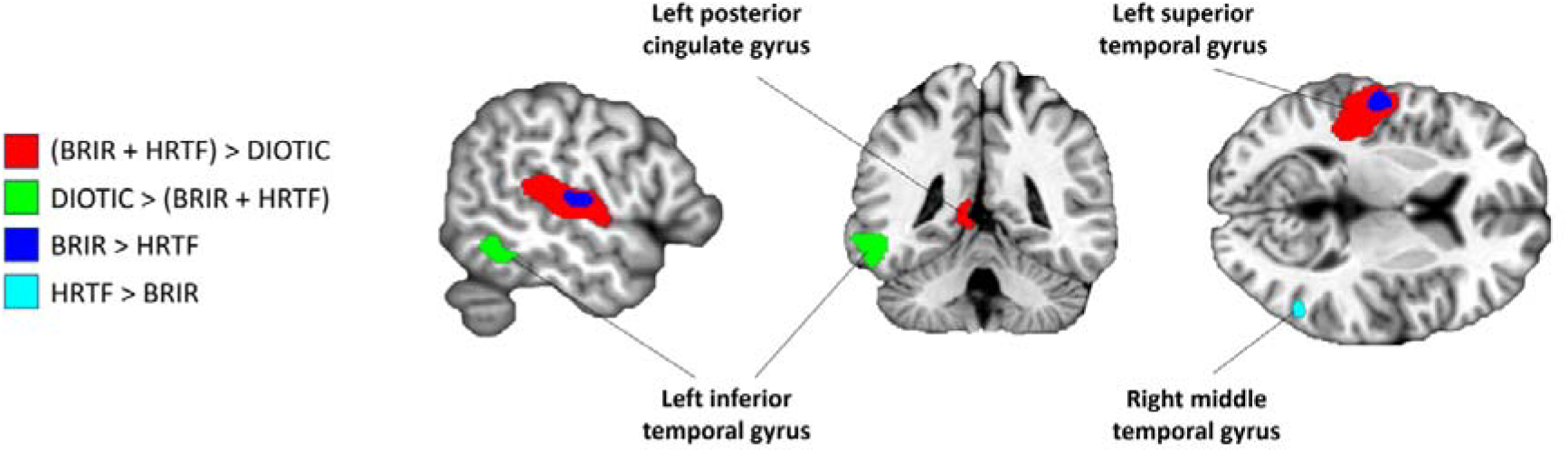
Summary of the contrast analyses of [BRIR + HRTF > diotic] (in *red*), [diotic > (HRTF + BRIR)] (in *green*), [BRIR > HRTF] (in *blue*) and [HRTF > BRIR] (in *turquoise*). The blue contrast overlaps with the red contrast in the left superior temporal gyrus. Images are overlaid on the Colin 27 anatomical template.

## 4. Discussion

Although our understanding of spatial sound localization has progressed, the neural underpinnings of externalization remain only partially understood. In this study, we investigated brain responses to three types of spatialized sound sources, BRIR, HRTF, and diotic stimuli, designed to modulate the perception of externalized and internalized sound sources. Our results revealed that externalized sounds elicited greater activity in the left superior temporal gyrus, including the PT, as well as in the cerebellum and the left posterior cingulate gyrus, compared to internalized sound sources. Conversely, internalized sounds were associated with greater relative activity in the left ITG compared to externalized sound sources.

The involvement of the left PT supports our hypothesis and corroborates previous findings implicating this region in auditory spatial processing (Hunter et al. 2003). As part of the postero-dorsal auditory path (van der Heijden et al. 2019), the PT contributes to acoustic spatial processing including distance perception (Kopčo et al. 2012), sound moving (Warren et al. 2002), and the integration of localization cues such as interaural time and level differences (Krumbholz et al. 2005b). The left-lateralized activation of the PT we observed may reflect the linguistic nature of our stimuli: vocalizations are processed similarly to speech, typically in the left hemisphere (Hunter et al. 2003), whereas non-vocal stimuli (e.g., noise) tend to evoke more bilateral responses (Callan et al. 2013). Additionally, the left-lateralized activation observed in the PT is consistent with the proposed third auditory stream, which supports the representation of meaningful sound objects in relation to their spatial position, distinct from both position-independent sound recognition and explicit sound localization (Clarke and Geiser 2015; Da Costa et al. 2018). Previous neuroimaging, auditory evoked potentials, and lesion studies have provided converging evidence for this stream and its left-hemisphere dominance (for a review see Clarke et al. 2024). From this perspective, the contrast between externalized and diotic sounds may reflect, at least in part, differences between position-dependent and position-independent representations of meaningful auditory stimuli. Our findings extend this literature by demonstrating left PT involvement with externalized vocalizations, providing further evidence for the contribution of this region to the processing of meaningful sounds in relation to their spatial position.

Our findings also resonate with studies in patients with auditory hallucinations, which report greater left PT activity among those who perceive their hallucinations as externally located, compared to those who perceive them as internal (Looijestijn et al., 2013). Taken together, these converging results suggest that the PT may play a crucial role in the attribution of spatial origin to auditory stimuli, both veridical and hallucinatory (Hunter, 2004), highlighting a research area that warrants further investigation.

The activation of the lobule VI of the cerebellum during externalized sound perception further supports this interpretation. This region has been implicated in reality monitoring processes that help differentiate internally generated experiences, such as imagination, from externally perceived ones. It also shows functional overlap with neural correlates of self-monitoring, the ability to distinguish self-generated actions or thoughts from those generated by others (Lavallé et al. 2023). Previous work has suggested that the cerebellum plays a role in sensory predictions of action and self-productions such as inner speech (Pinheiro et al. 2020), suggesting that its involvement in auditory externalization may reflect engagement of similar higher-order predictive and top-down mechanisms. In addition, given the role of the cerebellum in attention (Zhang et al. 2023), its activation further indicates that attentional mechanisms may contribute to the perception of externalized sounds. In line with this hypothesis, an electroencephalography study showed that BRIR-convolved sounds engage attention-related brain networks (Heine et al. 2021). The authors proposed that BRIR-convolved sounds enhance localization information, thereby increasing attentional engagement with auditory sources via top-down processes. This mechanism may enhance perceived externalization by directing attention toward external sound sources. Altogether, these findings suggest that auditory externalization may recruit a broader network involving not only sensory regions but also higher-level systems supporting prediction, attentional modulation, and source attribution (Heine et al. 2021; Lavallé et al. 2023).

The left posterior cingulate gyrus showed relative greater activity for externalized compared to internalized sound sources. This region, part of the default mode network, could be associated with internally-oriented cognition and tends to disengage during cognitively demanding tasks (Leech and Sharp 2014), which may be contradictory to our findings. However, alternative evidence suggests that the posterior cingulate may instead be involved in controlling the balance between internal and external attentional focus (Leech and Sharp 2014). Furthermore, the posterior cingulate has been implicated in somatosensory processing, spatial representation and visual dorsal stream (for a review, see Rolls 2019). Given its connectivity with the parietal cortex, this region may also participate in the dorsal auditory pathway, which could explain its relative increase in activity during the perception of externalized sounds compared to internalized ones. This aligns with the notion that accurate localization of sounds in space may require integrated body representation and spatial awareness.

Regarding internalized sound sources, our results revealed increased relative activity in the left ITG, extending to the middle temporal gyrus, suggesting potential involvement of the auditory spatial pathway. A recent study found that inner speech engages auditory processing areas even in the absence of external input (Stephane et al. 2021). The internal auditory experience of an inner voice could be spatially analogous to diotic sounds, which have no localization clues and yet activate the middle temporal gyrus. Additionally, we found greater activity in the right middle temporal gyrus in response to HRTF-convolved sounds, typically perceived as less externalized than BRIR-convolved sounds (Leclère et al. 2019). These findings corroborate the idea that internalized sounds may also engage spatial auditory networks.

Finally, while our findings replicated some previous results, we have also identified some discrepancies. For instance, the absence of significant activation in the inferior parietal lobe is surprising since its role in the auditory “where” pathway (Callan et al. 2013). This result could indicate that it is similarly activated by internalized and externalized sound sources (Stephane et al. 2021). Several methodological differences and limitations of our study could also contribute to this observation. For instance, the use of vocalizations instead of speech or noise could also explain this observation given that externalization is influenced by the acoustic characteristics of sounds (Heine et al. 2021). Future studies could therefore benefit from systematically comparing different types of stimuli (e.g., speech, non-speech vocalizations, environmental sounds) to assess the generalizability of the present findings. Another limitation is the use of non-individualized HRTF and BRIR stimuli. Some studies have reported improved externalization when individualized filters (derived from the listener’s own HRTFs) were used compared to generic ones. However, the added value of the individualization of the stimuli on externalization remains debated, as a substantial number of studies have found no clear advantage of individualized over non-individualized stimuli (for a review, see Best et al. 2020). In addition, although MRI provides great spatial resolution, it has a lower temporal resolution than electroencephalography and is susceptible to scanner noise. We mitigated noise-related effects by maintaining a consistent MRI background noise throughout the task to standardize the auditory environment.

To sum up, our study provides new insights into the neural underpinnings of auditory externalization. By employing an ecologically valid paradigm that simulates realistic sound sources, we identified key brain regions associated with the perception of externalized sound sources, including the superior temporal gyrus—particularly the left PT—the cerebellar lobule VI and posterior cingulate gyrus. In contrast, internalization was associated with increased activation in the left inferior and middle temporal gyri. Building on the present findings, future studies should aim to clarify how the neural mechanisms differentiating external and internal auditory experiences interface with higher-level cognitive processes, including source monitoring.

## Acknowledgements

The authors thank the CERMEP imaging center, in particular Franck LAMBERTON and Danielle IBARROLA for their help with the study design, the access to MRI, and data collection. We also thank all study participants.

## Statements & Declarations

### Author Contributions

Laure FIVEL: Conceptualization, Methodology, Software, Investigation, Formal analysis, Writing – original draft, Writing – review & editing, Visualization. Jérôme BRUNELIN: Conceptualization, Writing – review & editing. Gaëlle LEROUX: Methodology, Resources, Writing – review & editing. Frédéric HAESEBAERT: Conceptualization, Methodology, Writing – review & editing, Supervision, Funding acquisition. Marine MONDINO: Conceptualization, Methodology, Writing – review & editing, Visualisation, Supervision, Funding acquisition.

### Data Availability

The datasets generated during and analysed during the current study are not publicly available due to ethical restrictions on the sharing of individual-level data but aggregated group-level data may be available from the corresponding author on reasonable request.

