## Supplemental material for "How the Brain Processes Internal and External Sounds: An fMRI Investigation of Auditory Sound Externalization"

**Supplementary Material 1.** OptimizeX.

In each run, trials were randomized using OptimizeX (<http://www.bobspunt.com/easy-optimize-x/>) in Matlab 2020a with the following parameters for 1 run:

| **Parameters** | **Values** | **Details** |
| --- | --- | --- |
| TR | 1.5 | In seconds |
| High-Pass Filter cutoff (hpf) | 128 | In seconds |
| N Conditions (nconds) | 3 | BRIR, HRTF, diotic |
| N Trials per condition (ntrials) | 20 20 20 | 20 BRIR, 20 HRTF, 20 diotic |
| Trials from same condition occurring in a row (maxRep) | 3 | Maximum 3 sounds of the same condition can follow each other |
| Trial duration (trialDur) | 3 | The sound: 1s + the response: 2s |
| ISI (mean, min, max) | 3 2 4 | Inter Stimulus Interval in seconds |
| Optimization method of the ISI distribution (distISI) | Rayleigh | Choice between Rayleigh (default), chisquare and exponential |
| Rest interval to add to beginning of scan (restBegin) | 15 | In seconds |
| Rest interval to add at end of scan (restEnd) | 20 | In seconds |
| N of optimal designs to save (keep) | 6 |  |
| N of generations to test (ngen) | 100 |  |
| N of designs to include in each generation (gensize) | 500 |  |
| Max amount of time to run the program (maxtime) | 2 | In minutes |
| Contrasts | 2 | BRIR-DIOTIC; HRTF-DIOTIC |

We have launched OptimizeX with these parameters 6 times to obtain 6 different runs of 6 minutes. The results are displayed in the tables below. Then, we checked the expected results for the run in SPM12 to ensure the correct feasibility.

| **Run_1** | | | | | **Run_2** | | | | | **Run_3** | | | | |
| --- | --- | --- | --- | --- | --- | --- | --- | --- | --- | --- | --- | --- | --- | --- |
| Trial | Condition | Onset | Duration | ISI | Trial | Condition | Onset | Duration | ISI | Trial | Condition | Onset | Duration | ISI |
| 1 | 2 | 15.000 | 3 | 3.163 | 1 | 3 | 15.000 | 3 | 3.525 | 1 | 2 | 15.000 | 3 | 3.235 |
| 2 | 1 | 21.163 | 3 | 2.785 | 2 | 2 | 21.525 | 3 | 2.140 | 2 | 1 | 21.235 | 3 | 3.179 |
| 3 | 1 | 26.948 | 3 | 2.301 | 3 | 2 | 26.666 | 3 | 2.386 | 3 | 1 | 27.415 | 3 | 3.541 |
| 4 | 1 | 32.250 | 3 | 2.366 | 4 | 1 | 32.052 | 3 | 2.001 | 4 | 3 | 33.956 | 3 | 2.023 |
| 5 | 1 | 37.616 | 3 | 3.163 | 5 | 1 | 37.053 | 3 | 3.657 | 5 | 3 | 38.979 | 3 | 2.178 |
| 6 | 1 | 43.780 | 3 | 2.462 | 6 | 3 | 43.711 | 3 | 2.006 | 6 | 3 | 44.157 | 3 | 2.852 |
| 7 | 2 | 49.242 | 3 | 2.000 | 7 | 3 | 48.717 | 3 | 2.299 | 7 | 2 | 50.009 | 3 | 2.470 |
| 8 | 2 | 54.243 | 3 | 3.993 | 8 | 2 | 54.016 | 3 | 3.860 | 8 | 2 | 55.479 | 3 | 3.520 |
| 9 | 3 | 61.236 | 3 | 2.354 | 9 | 2 | 60.877 | 3 | 2.900 | 9 | 1 | 61.999 | 3 | 2.268 |
| 10 | 3 | 66.591 | 3 | 2.915 | 10 | 2 | 66.777 | 3 | 3.656 | 10 | 1 | 67.268 | 3 | 2.355 |
| 11 | 1 | 72.506 | 3 | 2.778 | 11 | 3 | 73.434 | 3 | 3.701 | 11 | 3 | 72.624 | 3 | 2.031 |
| 12 | 2 | 78.285 | 3 | 3.485 | 12 | 1 | 80.135 | 3 | 3.608 | 12 | 3 | 77.655 | 3 | 3.827 |
| 13 | 2 | 84.770 | 3 | 2.667 | 13 | 1 | 86.743 | 3 | 2.566 | 13 | 2 | 84.483 | 3 | 2.253 |
| 14 | 2 | 90.438 | 3 | 2.561 | 14 | 1 | 92.309 | 3 | 3.836 | 14 | 2 | 89.736 | 3 | 3.304 |
| 15 | 3 | 95.999 | 3 | 2.010 | 15 | 3 | 99.145 | 3 | 3.113 | 15 | 1 | 96.040 | 3 | 3.186 |
| 16 | 3 | 101.010 | 3 | 3.454 | 16 | 2 | 105.258 | 3 | 2.978 | 16 | 1 | 102.227 | 3 | 2.330 |
| 17 | 1 | 107.465 | 3 | 2.708 | 17 | 2 | 111.237 | 3 | 2.099 | 17 | 1 | 107.557 | 3 | 2.131 |
| 18 | 1 | 113.173 | 3 | 3.626 | 18 | 2 | 116.336 | 3 | 3.980 | 18 | 3 | 112.688 | 3 | 2.057 |
| 19 | 2 | 119.799 | 3 | 3.606 | 19 | 3 | 123.317 | 3 | 2.218 | 19 | 3 | 117.745 | 3 | 2.422 |
| 20 | 2 | 126.406 | 3 | 2.287 | 20 | 3 | 128.535 | 3 | 3.703 | 20 | 3 | 123.168 | 3 | 3.697 |
| 21 | 2 | 131.693 | 3 | 2.582 | 21 | 1 | 135.239 | 3 | 2.144 | 21 | 2 | 129.865 | 3 | 2.283 |
| 22 | 1 | 137.275 | 3 | 2.744 | 22 | 1 | 140.383 | 3 | 3.714 | 22 | 2 | 135.148 | 3 | 3.803 |
| 23 | 3 | 143.019 | 3 | 2.388 | 23 | 1 | 147.098 | 3 | 3.039 | 23 | 2 | 141.951 | 3 | 2.428 |
| 24 | 3 | 148.407 | 3 | 2.918 | 24 | 2 | 153.137 | 3 | 3.940 | 24 | 1 | 147.380 | 3 | 3.632 |
| 25 | 3 | 154.326 | 3 | 2.667 | 25 | 2 | 160.077 | 3 | 2.731 | 25 | 1 | 154.013 | 3 | 3.541 |
| 26 | 2 | 159.994 | 3 | 2.778 | 26 | 2 | 165.809 | 3 | 2.001 | 26 | 3 | 160.554 | 3 | 2.268 |
| 27 | 1 | 165.772 | 3 | 2.125 | 27 | 2 | 170.810 | 3 | 3.454 | 27 | 3 | 165.823 | 3 | 2.689 |
| 28 | 1 | 170.897 | 3 | 3.012 | 28 | 3 | 177.265 | 3 | 2.422 | 28 | 2 | 171.512 | 3 | 3.177 |
| 29 | 1 | 176.910 | 3 | 3.517 | 29 | 2 | 182.687 | 3 | 2.573 | 29 | 2 | 177.690 | 3 | 3.961 |
| 30 | 2 | 183.428 | 3 | 3.042 | 30 | 1 | 188.260 | 3 | 3.657 | 30 | 3 | 184.652 | 3 | 2.784 |
| 31 | 2 | 189.470 | 3 | 3.488 | 31 | 3 | 194.918 | 3 | 2.273 | 31 | 3 | 190.436 | 3 | 3.446 |
| 32 | 3 | 195.959 | 3 | 2.078 | 32 | 3 | 200.191 | 3 | 2.270 | 32 | 2 | 196.882 | 3 | 2.283 |
| 33 | 3 | 201.037 | 3 | 3.913 | 33 | 3 | 205.461 | 3 | 3.742 | 33 | 2 | 202.166 | 3 | 3.446 |
| 34 | 1 | 207.951 | 3 | 2.358 | 34 | 1 | 212.204 | 3 | 3.836 | 34 | 1 | 208.613 | 3 | 3.199 |
| 35 | 2 | 213.309 | 3 | 3.488 | 35 | 2 | 219.041 | 3 | 2.512 | 35 | 1 | 214.813 | 3 | 3.024 |
| 36 | 2 | 219.798 | 3 | 3.398 | 36 | 1 | 224.554 | 3 | 3.682 | 36 | 2 | 220.837 | 3 | 3.349 |
| 37 | 3 | 226.196 | 3 | 2.744 | 37 | 1 | 231.236 | 3 | 3.809 | 37 | 1 | 227.186 | 3 | 3.519 |
| 38 | 3 | 231.940 | 3 | 3.906 | 38 | 3 | 238.045 | 3 | 2.617 | 38 | 1 | 233.706 | 3 | 2.281 |
| 39 | 1 | 238.847 | 3 | 2.785 | 39 | 3 | 243.663 | 3 | 3.502 | 39 | 1 | 238.987 | 3 | 3.433 |
| 40 | 2 | 244.633 | 3 | 3.158 | 40 | 2 | 250.166 | 3 | 2.035 | 40 | 2 | 245.420 | 3 | 2.495 |
| 41 | 3 | 250.791 | 3 | 2.582 | 41 | 2 | 255.201 | 3 | 2.270 | 41 | 2 | 250.916 | 3 | 2.962 |
| 42 | 3 | 256.373 | 3 | 2.089 | 42 | 1 | 260.471 | 3 | 2.743 | 42 | 3 | 256.879 | 3 | 2.574 |
| 43 | 3 | 261.46 | 3 | 2.582 | 43 | 1 | 266.215 | 3 | 3.960 | 43 | 3 | 262.453 | 3 | 3.987 |
| 44 | 3 | 267.045 | 3 | 2.166 | 44 | 3 | 273.175 | 3 | 2.434 | 44 | 3 | 269.440 | 3 | 3.755 |
| 45 | 2 | 272.211 | 3 | 3.632 | 45 | 3 | 278.609 | 3 | 2.311 | 45 | 2 | 276.196 | 3 | 2.137 |
| 46 | 2 | 278.844 | 3 | 3.873 | 46 | 2 | 283.920 | 3 | 2.612 | 46 | 2 | 281.333 | 3 | 3.756 |
| 47 | 1 | 285.717 | 3 | 2.276 | 47 | 1 | 289.533 | 3 | 3.232 | 47 | 3 | 288.090 | 3 | 3.252 |
| 48 | 1 | 290.993 | 3 | 2.556 | 48 | 1 | 295.766 | 3 | 2.385 | 48 | 3 | 294.342 | 3 | 2.962 |
| 49 | 3 | 296.549 | 3 | 2.558 | 49 | 1 | 301.151 | 3 | 3.381 | 49 | 2 | 300.305 | 3 | 3.278 |
| 50 | 3 | 302.108 | 3 | 3.958 | 50 | 3 | 307.532 | 3 | 2.461 | 50 | 1 | 306.583 | 3 | 2.468 |
| 51 | 1 | 309.067 | 3 | 2.934 | 51 | 3 | 312.993 | 3 | 2.849 | 51 | 1 | 312.051 | 3 | 3.311 |
| 52 | 1 | 315.001 | 3 | 2.982 | 52 | 3 | 318.843 | 3 | 2.965 | 52 | 1 | 318.363 | 3 | 3.070 |
| 53 | 1 | 320.983 | 3 | 3.776 | 53 | 2 | 324.808 | 3 | 3.125 | 53 | 2 | 324.433 | 3 | 2.848 |
| 54 | 2 | 327.760 | 3 | 3.888 | 54 | 1 | 330.934 | 3 | 2.298 | 54 | 1 | 330.281 | 3 | 3.379 |
| 55 | 3 | 334.648 | 3 | 2.240 | 55 | 1 | 336.233 | 3 | 2.400 | 55 | 3 | 336.661 | 3 | 3.359 |
| 56 | 3 | 339.888 | 3 | 3.738 | 56 | 2 | 341.634 | 3 | 2.450 | 56 | 3 | 343.021 | 3 | 3.208 |
| 57 | 1 | 346.627 | 3 | 2.366 | 57 | 2 | 347.084 | 3 | 3.741 | 57 | 1 | 349.230 | 3 | 3.301 |
| 58 | 2 | 351.993 | 3 | 2.574 | 58 | 3 | 353.826 | 3 | 3.115 | 58 | 2 | 355.531 | 3 | 3.595 |
| 59 | 2 | 357.568 | 3 | 3.915 | 59 | 3 | 359.942 | 3 | 3.297 | 59 | 3 | 362.127 | 3 | 3.354 |
| 60 | 3 | 364.484 | 3 | 20.000 | 60 | 1 | 366.240 | 3 | 20.000 | 60 | 1 | 368.481 | 3 | 20.000 |

Supp. table 1. Onsets and duration for each trial for runs 1 to run 3. Condition 1 = BRIR, 2=HRTF, 3 = diotic.

Supp. table 2. From run 4 to run 6. Condition 1 = BRIR, 2=HRTF, 3 = diotic.

| **Run_4** | | | | | **Run_5** | | | | | **Run_6** | | | | |
| --- | --- | --- | --- | --- | --- | --- | --- | --- | --- | --- | --- | --- | --- | --- |
| Trial | Condition | Onset | Duration | ISI | Trial | Condition | Onset | Duration | ISI | Trial | Condition | Onset | Duration | ISI |
| 1 | 2 | 15.000 | 3 | 2.186 | 1 | 3 | 15.000 | 3 | 2.962 | 1 | 3 | 15.000 | 3 | 3.358 |
| 2 | 1 | 20.186 | 3 | 3.304 | 2 | 1 | 20.962 | 3 | 2.631 | 2 | 1 | 21.358 | 3 | 3.114 |
| 3 | 3 | 26.490 | 3 | 3.338 | 3 | 1 | 26.594 | 3 | 2.383 | 3 | 2 | 27.472 | 3 | 2.451 |
| 4 | 1 | 32.829 | 3 | 2.622 | 4 | 1 | 31.978 | 3 | 2.731 | 4 | 3 | 32.924 | 3 | 2.789 |
| 5 | 1 | 38.451 | 3 | 3.375 | 5 | 2 | 37.709 | 3 | 3.500 | 5 | 3 | 38.713 | 3 | 3.968 |
| 6 | 3 | 44.827 | 3 | 2.219 | 6 | 3 | 44.210 | 3 | 2.870 | 6 | 1 | 45.682 | 3 | 2.458 |
| 7 | 3 | 50.047 | 3 | 3.597 | 7 | 1 | 50.080 | 3 | 2.668 | 7 | 1 | 51.140 | 3 | 3.136 |
| 8 | 1 | 56.644 | 3 | 3.140 | 8 | 3 | 55.748 | 3 | 3.387 | 8 | 1 | 57.277 | 3 | 3.082 |
| 9 | 2 | 62.785 | 3 | 3.647 | 9 | 2 | 62.136 | 3 | 2.685 | 9 | 3 | 63.359 | 3 | 2.050 |
| 10 | 1 | 69.433 | 3 | 3.373 | 10 | 1 | 67.821 | 3 | 2.525 | 10 | 3 | 68.409 | 3 | 3.974 |
| 11 | 3 | 75.806 | 3 | 2.697 | 11 | 2 | 73.346 | 3 | 3.464 | 11 | 2 | 75.383 | 3 | 2.217 |
| 12 | 3 | 81.504 | 3 | 2.062 | 12 | 2 | 79.811 | 3 | 2.646 | 12 | 2 | 80.600 | 3 | 2.885 |
| 13 | 1 | 86.566 | 3 | 3.251 | 13 | 1 | 85.458 | 3 | 3.294 | 13 | 2 | 86.486 | 3 | 3.465 |
| 14 | 2 | 92.818 | 3 | 2.966 | 14 | 3 | 91.753 | 3 | 3.634 | 14 | 1 | 92.952 | 3 | 2.597 |
| 15 | 2 | 98.784 | 3 | 2.862 | 15 | 3 | 98.387 | 3 | 2.712 | 15 | 1 | 98.549 | 3 | 2.710 |
| 16 | 2 | 104.646 | 3 | 3.443 | 16 | 1 | 104.099 | 3 | 2.883 | 16 | 3 | 104.260 | 3 | 2.987 |
| 17 | 1 | 111.089 | 3 | 2.617 | 17 | 1 | 109.982 | 3 | 2.368 | 17 | 2 | 110.247 | 3 | 2.165 |
| 18 | 1 | 116.707 | 3 | 3.918 | 18 | 2 | 115.350 | 3 | 2.972 | 18 | 1 | 115.412 | 3 | 2.191 |
| 19 | 3 | 123.626 | 3 | 2.581 | 19 | 3 | 121.323 | 3 | 2.583 | 19 | 3 | 120.603 | 3 | 2.818 |
| 20 | 3 | 129.208 | 3 | 3.947 | 20 | 3 | 126.906 | 3 | 2.008 | 20 | 2 | 126.422 | 3 | 3.385 |
| 21 | 2 | 136.155 | 3 | 3.373 | 21 | 3 | 131.914 | 3 | 2.008 | 21 | 1 | 132.807 | 3 | 2.344 |
| 22 | 2 | 142.529 | 3 | 2.530 | 22 | 2 | 136.922 | 3 | 2.055 | 22 | 1 | 138.151 | 3 | 3.209 |
| 23 | 1 | 148.059 | 3 | 2.752 | 23 | 1 | 141.977 | 3 | 2.253 | 23 | 3 | 144.361 | 3 | 2.519 |
| 24 | 1 | 153.812 | 3 | 3.802 | 24 | 1 | 147.231 | 3 | 3.211 | 24 | 3 | 149.881 | 3 | 3.677 |
| 25 | 3 | 160.614 | 3 | 2.278 | 25 | 2 | 153.443 | 3 | 3.482 | 25 | 1 | 156.558 | 3 | 2.458 |
| 26 | 3 | 165.893 | 3 | 2.723 | 26 | 2 | 159.925 | 3 | 3.387 | 26 | 1 | 162.017 | 3 | 2.030 |
| 27 | 2 | 171.616 | 3 | 2.218 | 27 | 3 | 166.312 | 3 | 2.486 | 27 | 2 | 167.048 | 3 | 2.707 |
| 28 | 2 | 176.835 | 3 | 3.483 | 28 | 3 | 171.799 | 3 | 3.928 | 28 | 2 | 172.756 | 3 | 3.772 |
| 29 | 1 | 183.318 | 3 | 3.552 | 29 | 2 | 178.728 | 3 | 2.530 | 29 | 2 | 179.528 | 3 | 3.564 |
| 30 | 1 | 189.870 | 3 | 3.757 | 30 | 1 | 184.258 | 3 | 2.814 | 30 | 3 | 186.093 | 3 | 2.719 |
| 31 | 2 | 196.627 | 3 | 3.008 | 31 | 1 | 190.072 | 3 | 3.103 | 31 | 1 | 191.813 | 3 | 2.170 |
| 32 | 3 | 202.635 | 3 | 2.089 | 32 | 1 | 196.175 | 3 | 2.993 | 32 | 1 | 196.983 | 3 | 3.900 |
| 33 | 3 | 207.725 | 3 | 3.135 | 33 | 3 | 202.169 | 3 | 2.477 | 33 | 3 | 203.884 | 3 | 3.114 |
| 34 | 1 | 213.860 | 3 | 3.098 | 34 | 2 | 207.647 | 3 | 3.941 | 34 | 3 | 209.998 | 3 | 2.093 |
| 35 | 2 | 219.959 | 3 | 2.058 | 35 | 1 | 214.589 | 3 | 3.278 | 35 | 3 | 215.091 | 3 | 2.832 |
| 36 | 3 | 225.018 | 3 | 3.986 | 36 | 3 | 220.868 | 3 | 2.719 | 36 | 2 | 220.924 | 3 | 3.469 |
| 37 | 1 | 232.004 | 3 | 2.968 | 37 | 3 | 226.588 | 3 | 3.310 | 37 | 1 | 227.393 | 3 | 2.619 |
| 38 | 2 | 237.973 | 3 | 3.645 | 38 | 2 | 232.899 | 3 | 3.755 | 38 | 1 | 233.013 | 3 | 2.033 |
| 39 | 3 | 244.618 | 3 | 3.553 | 39 | 2 | 239.655 | 3 | 2.180 | 39 | 1 | 238.046 | 3 | 2.307 |
| 40 | 3 | 251.171 | 3 | 3.458 | 40 | 2 | 244.835 | 3 | 3.725 | 40 | 3 | 243.353 | 3 | 3.564 |
| 41 | 2 | 257.630 | 3 | 2.690 | 41 | 1 | 251.560 | 3 | 2.919 | 41 | 2 | 249.918 | 3 | 2.917 |
| 42 | 2 | 263.320 | 3 | 2.723 | 42 | 3 | 257.479 | 3 | 2.357 | 42 | 2 | 255.835 | 3 | 2.675 |
| 43 | 1 | 269.044 | 3 | 2.386 | 43 | 3 | 262.837 | 3 | 3.549 | 43 | 2 | 261.511 | 3 | 2.340 |
| 44 | 1 | 274.430 | 3 | 3.986 | 44 | 2 | 269.386 | 3 | 3.374 | 44 | 3 | 266.852 | 3 | 2.101 |
| 45 | 3 | 281.417 | 3 | 3.297 | 45 | 1 | 275.760 | 3 | 2.327 | 45 | 3 | 271.953 | 3 | 3.453 |
| 46 | 1 | 287.714 | 3 | 2.916 | 46 | 1 | 281.088 | 3 | 2.684 | 46 | 1 | 278.407 | 3 | 2.152 |
| 47 | 2 | 293.631 | 3 | 2.261 | 47 | 3 | 286.772 | 3 | 2.786 | 47 | 1 | 283.559 | 3 | 3.260 |
| 48 | 2 | 298.892 | 3 | 3.135 | 48 | 3 | 292.558 | 3 | 2.651 | 48 | 3 | 289.819 | 3 | 3.174 |
| 49 | 3 | 305.027 | 3 | 2.966 | 49 | 2 | 298.210 | 3 | 2.565 | 49 | 2 | 295.993 | 3 | 2.762 |
| 50 | 3 | 310.993 | 3 | 3.818 | 50 | 2 | 303.776 | 3 | 3.651 | 50 | 1 | 301.756 | 3 | 3.666 |
| 51 | 2 | 317.812 | 3 | 3.648 | 51 | 1 | 310.428 | 3 | 3.277 | 51 | 2 | 308.423 | 3 | 2.211 |
| 52 | 1 | 324.461 | 3 | 3.288 | 52 | 1 | 316.705 | 3 | 3.761 | 52 | 2 | 313.634 | 3 | 3.924 |
| 53 | 3 | 330.749 | 3 | 2.248 | 53 | 3 | 323.467 | 3 | 3.279 | 53 | 3 | 320.558 | 3 | 2.302 |
| 54 | 3 | 335.997 | 3 | 3.158 | 54 | 2 | 329.747 | 3 | 2.604 | 54 | 3 | 325.861 | 3 | 3.752 |
| 55 | 2 | 342.156 | 3 | 2.747 | 55 | 2 | 335.352 | 3 | 2.120 | 55 | 2 | 332.613 | 3 | 2.917 |
| 56 | 2 | 347.904 | 3 | 3.320 | 56 | 2 | 340.472 | 3 | 3.661 | 56 | 2 | 338.530 | 3 | 2.549 |
| 57 | 2 | 354.224 | 3 | 2.054 | 57 | 3 | 347.134 | 3 | 2.434 | 57 | 3 | 344.079 | 3 | 3.752 |
| 58 | 1 | 359.279 | 3 | 2.464 | 58 | 3 | 352.568 | 3 | 3.459 | 58 | 1 | 350.832 | 3 | 3.516 |
| 59 | 1 | 364.743 | 3 | 3.691 | 59 | 2 | 359.028 | 3 | 3.588 | 59 | 2 | 357.348 | 3 | 3.948 |
| 60 | 3 | 371.435 | 3 | 20.000 | 60 | 1 | 365.617 | 3 | 20.000 | 60 | 2 | 364.297 | 3 | 20.000 |

**Supplementary Material 2.** fMRI preprocessing

Results included in this manuscript come from preprocessing performed using fMRIPrep 23.2.0 (Esteban et al. (2019); Esteban et al. (2018); RRID:SCR_016216), which is based on Nipype 1.8.6 (Gorgolewski et al. (2011); Gorgolewski et al. (2018); RRID:SCR_002502).

**Preprocessing of B0 inhomogeneity mappings**

A total of 1 fieldmap was found available within the input BIDS structure for this particular subject. A B0 nonuniformity map (or fieldmap) was estimated from the phase-drift map(s) measure with two consecutive GRE (gradient-recalled echo) acquisitions. The corresponding phase-map(s) were phase-unwrapped with prelude (FSL None).

**Anatomical data preprocessing**

A total of 1 T1-weighted (T1w) image was found within the input BIDS dataset. The T1w image was corrected for intensity non-uniformity (INU) with N4BiasFieldCorrection (Tustison et al. 2010), distributed with ANTs 2.5.0 (Avants et al. 2008, RRID:SCR_004757), and used as T1w-reference throughout the workflow. The T1w-reference was then skull-stripped with a Nipype implementation of the antsBrainExtraction.sh workflow (from ANTs), using OASIS30ANTs as target template. Brain tissue segmentation of cerebrospinal fluid (CSF), white-matter (WM) and gray-matter (GM) was performed on the brain-extracted T1w using fast (FSL (version unknown), RRID:SCR_002823, Zhang, Brady, and Smith 2001). Brain surfaces were reconstructed using recon-all (FreeSurfer 7.3.2, RRID:SCR_001847, Dale, Fischl, and Sereno 1999), and the brain mask estimated previously was refined with a custom variation of the method to reconcile ANTs-derived and FreeSurfer-derived segmentations of the cortical gray-matter of Mindboggle (RRID:SCR_002438, Klein et al. 2017). A T2-weighted image was used to improve pial surface refinement. Brain surfaces were reconstructed using recon-all (FreeSurfer 7.3.2, RRID:SCR_001847, Dale, Fischl, and Sereno 1999), and the brain mask estimated previously was refined with a custom variation of the method to reconcile ANTs-derived and FreeSurfer-derived segmentations of the cortical gray-matter of Mindboggle (RRID:SCR_002438, Klein et al. 2017). Volume-based spatial normalization to one standard space (MNI152NLin2009cAsym) was performed through nonlinear registration with antsRegistration (ANTs 2.5.0), using brain-extracted versions of both T1w reference and the T1w template. The following template was selected for spatial normalization and accessed with TemplateFlow (23.1.0, Ciric et al. 2022): ICBM 152 Nonlinear Asymmetrical template version 2009c [Fonov et al. (2009), RRID:SCR_008796; TemplateFlow ID: MNI152NLin2009cAsym].

**Functional data preprocessing**

For each of the 6 BOLD runs found per subject (across all tasks and sessions), the following preprocessing was performed. First, a reference volume was generated, using a custom methodology of fMRIPrep, for use in head motion correction. Head-motion parameters with respect to the BOLD reference (transformation matrices, and six corresponding rotation and translation parameters) are estimated before any spatiotemporal filtering using mcflirt (FSL, Jenkinson et al. 2002). The estimated fieldmap was then aligned with rigid-registration to the target EPI (echo-planar imaging) reference run. The field coefficients were mapped on to the reference EPI using the transform. The BOLD reference was then co-registered to the T1w reference using bbregister (FreeSurfer) which implements boundary-based registration (Greve and Fischl 2009). Co-registration was configured with six degrees of freedom. Several confounding time-series were calculated based on the preprocessed BOLD: framewise displacement (FD), DVARS and three region-wise global signals. FD was computed using two formulations following Power (absolute sum of relative motions, Power et al. (2014)) and Jenkinson (relative root mean square displacement between affines, Jenkinson et al. (2002)). FD and DVARS are calculated for each functional run, both using their implementations in Nipype (following the definitions by Power et al. 2014). The three global signals are extracted within the CSF, the WM, and the whole-brain masks. Additionally, a set of physiological regressors were extracted to allow for component-based noise correction (CompCor, Behzadi et al. 2007). Principal components are estimated after high-pass filtering the preprocessed BOLD time-series (using a discrete cosine filter with 128s cut-off) for the two CompCor variants: temporal (tCompCor) and anatomical (aCompCor). tCompCor components are then calculated from the top 2% variable voxels within the brain mask. For aCompCor, three probabilistic masks (CSF, WM and combined CSF+WM) are generated in anatomical space. The implementation differs from that of Behzadi et al. in that instead of eroding the masks by 2 pixels on BOLD space, a mask of pixels that likely contain a volume fraction of GM is subtracted from the aCompCor masks. This mask is obtained by dilating a GM mask extracted from the FreeSurfer’s aseg segmentation, and it ensures components are not extracted from voxels containing a minimal fraction of GM. Finally, these masks are resampled into BOLD space and binarized by thresholding at 0.99 (as in the original implementation). Components are also calculated separately within the WM and CSF masks. For each CompCor decomposition, the k components with the largest singular values are retained, such that the retained components’ time series are sufficient to explain 50 percent of variance across the nuisance mask (CSF, WM, combined, or temporal). The remaining components are dropped from consideration. The head-motion estimates calculated in the correction step were also placed within the corresponding confounds file. The confound time series derived from head motion estimates and global signals were expanded with the inclusion of temporal derivatives and quadratic terms for each (Satterthwaite et al. 2013). Frames that exceeded a threshold of 0.5 mm FD or 1.5 standardized DVARS were annotated as motion outliers. Additional nuisance timeseries are calculated by means of principal components analysis of the signal found within a thin band (crown) of voxels around the edge of the brain, as proposed by (Patriat, Reynolds, and Birn 2017). All resamplings can be performed with a single interpolation step by composing all the pertinent transformations (i.e. head-motion transform matrices, susceptibility distortion correction when available, and co-registrations to anatomical and output spaces). Gridded (volumetric) resamplings were performed using nitransforms, configured with cubic B-spline interpolation.

Many internal operations of fMRIPrep use Nilearn 0.10.2 (Abraham et al. 2014, RRID:SCR_001362), mostly within the functional processing workflow. For more details of the pipeline, see the section corresponding to workflows in fMRIPrep’s documentation.

**Copyright Waiver**

The above boilerplate text was automatically generated by fMRIPrep with the express intention that users should copy and paste this text into their manuscripts unchanged. It is released under the CC0 license.
